# A nickase Cas9 gene-drive system promotes super-Mendelian inheritance in *Drosophila*

**DOI:** 10.1101/2021.12.01.470847

**Authors:** Víctor López Del Amo, Sara Sanz Juste, Valentino M. Gantz

## Abstract

CRISPR-based gene-drive systems have been proposed for managing insect populations, including disease-transmitting mosquitoes, due to their ability to bias their inheritance towards super-Mendelian rates (>50%). Current technologies employ a Cas9 that introduces DNA double-strand breaks into the opposing wildtype allele to replace it with a copy of the gene drive allele via DNA homology-directed repair. Yet, the use of different Cas9s versions is unexplored, and alternative approaches could increase the available toolkit for gene-drive designs. Here, we report a novel gene-drive approach that relies on Cas9 nickases that generate staggered paired nicks in DNA to propagate the engineered gene-drive cassette. We show that generating 5’ overhangs in the system yields efficient allelic conversion. The nickase gene-drive arrangement produces large, stereotyped deletions that are advantageous for targeting essential genes. Indeed, our nickase approach should expand the repertoire for gene-drive designs aimed at applications in mosquitoes and beyond.

## INTRODUCTION

CRISPR gene-drive systems have emerged as a promising tool for disseminating engineered traits into wild populations to control disease transmission. This rapid dissemination is possible due to the ability of these systems to surpass Mendel’s First Law of gene segregation, which dictates that an allele has a 50% probability of being passed to the next generation; in fact, gene drives can reach up to 100% inheritance of a desired gene. A proof-of-concept system was first implemented in flies (Gantz and Bier, 2015) and was applied to different mosquitoes such as *Anopheles* or *Aedes* under laboratory conditions to fight vector-borne diseases (Adolfi et al., 2020; Gantz et al., 2015; Hammond et al., 2016; Kyrou et al., 2018; Li et al., 2020; Simoni et al., 2020). CRISPR-based gene drives consist of a three-component transgene: (i) Cas9, a DNA nuclease that produces DNA double-strand breaks; (ii) a guide RNA (gRNA) that directs Cas9 to cleave the DNA at a predetermined site; and (iii) two homology arms flanking the Cas9/gRNA components that perfectly match both sides of the cut site to promote homology-directed repair (HDR). When a gene-drive individual mates with a wildtype, the encoded Cas9 from the engineered gene drive cuts the wildtype allele in the germline, which is replaced by HDR using the intact gene-drive chromosome as a repair template. With the gene drive present on both alleles (i.e., homozygous), this process produces a super-Mendelian inheritance (>50%) of the engineered cassette to spread new traits through a population.

Current gene-drive methods employ a Cas9 that introduces DNA double-strand breaks (Adolfi et al., 2020; Bier, 2021; Gantz et al., 2015; Hammond et al., 2016; Kyrou et al., 2018; Li et al., 2020; Simoni et al., 2020). However, we lack alternative Cas9-based strategies that could enlarge the available toolkit for gene-drive designs while potentially bringing advantages to improve the current ones. In fact, mutant versions of Cas9 that only generate nicks should also be amenable. Wildtype Cas9 contains two endonuclease domains (HNH and RuvC-like domains) that can introduce DNA double-strand breaks, where each cleaves one strand of the DNA double-helix. By mutating critical residues in the nuclease domains, two nickase versions of Cas9 can be generated by mutating critical residues in the nuclease domains: i) nCas9-D10A (nD10A) contains an inactivated RuvC domain and only cuts the target strand where the gRNA is bound, and ii) nCas9-H840A (nH840A) contains an inactivated HNH and so only cuts the non-target strand (Jinek et al., 2012).

Importantly, the nickase versions of Cas9 already demonstrated their efficiency for genome editing. nD10A has been used to generate paired DNA nicks and efficiently disrupts genes in *Drosophila* and cell culture (Gopalappa et al., 2018; Port et al., 2014). Nicks induced by nD10A promoted higher HDR rates than nH840A, where HDR was almost undetectable (Bothmer et al., 2017; Hyodo et al., 2020; Mali et al., 2013; Wang et al., 2021, 2018). Furthermore, nD10A can boost specificity while reducing off-target effects since it requires two gRNAs that target complementary DNA strands to either disrupt a gene function or trigger HDR. When the same pair of gRNAs were combined with wildtype Cas9, undesired off-target effects at an unspecific genomic region were generated, which does not occur with nD10A (Ran et al., 2013).

While nickase Cas9 (nCas9) versions have been successfully utilized *in vitro*, to the best of our knowledge, none paired-gRNAs’ nickase-based approaches have demonstrated efficient HDR in the germline of a living organism. We reasoned that a nCas9-based gene drive might be applicable for population engineering to bring potential advantages. For example, DNA nicks are involved in important biological processes such as DNA replication and are typically repaired efficiently (Caldecott, 2008; Chafin et al., 2000; Reyes et al., 2021; Wang and Hays, 2007). Therefore, if paired nicks do not occur simultaneously, single nicks should restore the original wildtype sequence, reducing the formation of mutations or resistant alleles at the target site for further gene-drive conversion. Additionally, DNA nicks follow distinct DNA repair pathways compared to DNA double-strand breaks that are introduced by traditional gene drives (Vriend and Krawczyk, 2017), and the intrinsically offset distance between paired nicks in a gene-drive setting could potentially favor the formation of specific mutations to improve gene-drive efficiency in certain applications.

Thus, we envisioned that simultaneous paired nicks targeting two adjacent DNA regions should generate a staggered double-strand break followed by DNA repair by HDR to promote super-Mendelian inheritance of an engineered gene-drive construct. Herein, we developed a nCas9-based gene-drive system promoting super-Mendelian inheritance in *Drosophila melanogaster* as a proof-of-concept and showed that nD10A and nH840A can promote efficient HDR in the germline. Interestingly, we showed that super-Mendelian inheritance rates can be achieved only when the gene-drive design generated 5’ overhangs. We also showed that nH840A produces larger deletions compared to nD10A when the allelic conversion fails, which can potentially be employed to ensure gene-drive spread when targeting essential genes while providing a rescue for the disrupted gene as part of the driven cassette. Overall, this work expands the technology and applicability of CRISPR gene-drive systems for genetic engineering of wild populations.

## RESULTS

### Nickase gene-drive system can be tailored to induce 5’ or 3’ overhangs

To design a nickase-based gene drive, we used a gRNA-only split-drive system (i.e., Copycat) that consists of two separate components: (i) a transgenic fly carrying a static Cas9 transgene, which is inherited in a Mendelian fashion, and (ii) an engineered animal carrying a CopyCat cassette formed by a gRNA gene that is flanked by two homology arms (Gantz and Bier, 2016). Once the two components are genetically crossed, traditional CopyCat gene drives rely on DNA double-strand breaks produced by a single gRNA, which targets the same sequence on the wildtype allele where it is inserted, to propagate the synthetic cassette by HDR (Champer et al., 2019; Gantz and Bier, 2016; López Del Amo et al., 2020a; Xu et al., 2017) (**Fig.1a**). In contrast, our nickase-based gene-drive system requires two gRNAs since this modified Cas9 introduces nicks instead of double-strand breaks. In this case, the gRNA pair will produce two independent cleavage events on each of the complementary DNA strands for gene drive propagation by HDR, emulating traditional gene drives (**Fig.1b**).

To build a gene drive system based on a nCas9, we generated two transgenic lines containing a DsRed marker and carrying either the nD10A or nH840A versions, which cut the target strand (bound to the gRNA) or the non-target strand, respectively. Additionally, we employed a validated wildtype Cas9 line, which introduces DNA double-strand breaks (López Del Amo et al., 2020b) as a positive control (**Fig. 1c**). We inserted all Cas9 transgenes into the *yellow* locus and expressed them with the same *vasa* germline promoter. Separately, we built two Copycat gene-drive constructs that we inserted into the *white* gene to produce two distinct gRNAs. To ensure perfect homology and a proper HDR process, the two homology arms included as part of the CopyCat elements match each cut site of the gRNA pair. We used a GFP marker to track the inheritance of these transgenes. Both Copycat lines share the *w2*-gRNA, which we previously validated (López Del Amo et al., 2020a, 2020b), and were combined with either the *w8*-gRNA or the *w9*-gRNA as the second gRNA (**Fig. 1d**). Additionally, PAM DNA sequences are crucial for target location recognition (Jinek et al., 2012), and these constructs present different PAM orientations depending on the gene-drive element. The Copycat transgenic line containing *w2,w8*-gRNAs pairs (CC(*w2,w8*)) have PAMs that are facing in opposite directions (i.e., a PAM-out orientation). In contrast, the transgenic strain carrying the *w2,w9*-gRNAs pairs (CC(*w2,w9*)) have PAMs that are facing each other (i.e., a PAM-in orientation). All three gRNAs are located within a ∼100 base pair DNA window, and the paired gRNA cut sites are separated by approximately 50 nucleotides (**Fig. 1d**).

**Fig.1.**
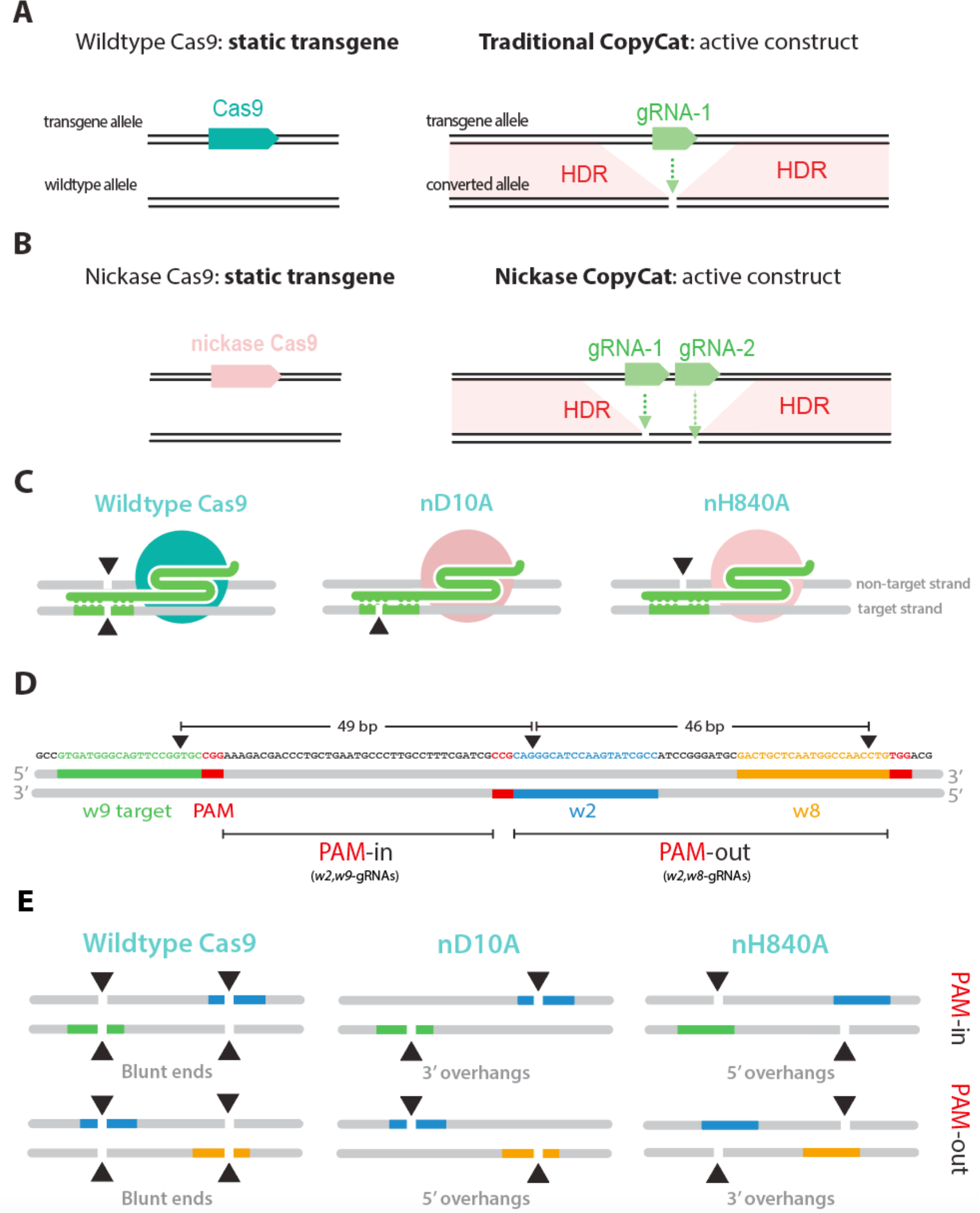
A nickase-based gene drive system promotes different overhang patterns. **a**. Schematic diagram of a traditional CopyCat gene-drive system. When combined with a Cas9 source, the gRNA cassette replaces the wildtype allele (converted allele) by DNA double-strand break and subsequent homology directed-repair (HDR). **b**. A nickase Cas9 source is combined with a Copycat containing two gRNAs targeting each complementary strand of the wildtype allele to spread the paired gRNA cassette by HDR. **c**. Wildtype Cas9 cuts both DNA strands, nD10A cuts the target strand where the gRNA is bound, and nH840 cuts the non-target strand. **d**. Sequence and design of the paired gRNAs in both PAM-out and PAM-in orientation. Paired gRNAs are located ∼50 nucleotides apart. The depicted gRNAs bind to the opposite strand when produced by complementarity. Red boxes indicate the PAM sequences (not included in the gRNA) that are crucial for DNA recognition. The black triangles denote the different cut sites associated with each gRNA **e**. Wildtype Cas9 introduces blunt ends when combined with either of the CopyCat elements. nD10A and nH840, combined with paired gRNAs binding to specific DNA strands, can generate 5’ or 3’ overhangs as they target different strands (target and non-target strands, respectively).

We combined the nickases with the two CopyCat transgenic lines to create four schemes to test the nickase gene-drive system: i) nD10A with the CC(*w2,w9*) (PAM-in) to generate 3’ overhangs; ii) nD10A with the CC(*w2,w8*) (PAM-out) to generate 5’ overhangs; iii) nH840A with CC(*w2,w9*) (PAM-in) to generate 5’ overhangs and iv) nH840A with the CC(*w2,w8*) (PAM-out) transgenic line to generate 3’ overhangs. Importantly, combining either of the CopyCat lines with wildtype Cas9 produces similar blunt ends in both situations (**Fig. 1e**).

### Cas9 nickases promote super-Mendelian inheritance of the CopyCat elements

To evaluate the efficiency of our CopyCat elements, we used the same genetic cross scheme in all cases. First, we combined male flies containing the Cas9 source (wildtype Cas9, nD10A, or nH840A) with the CopyCat lines to obtain F_1_ trans-heterozygous animals carrying both transgenes. We then crossed these F_1_ females to a white mutant line to evaluate biased inheritance in their F_2_ progeny. If the Copycat is inactive, we should observe 50% inheritance of the GFP marker, which is integrated within the engineered cassette as mentioned above. If the CopyCat construct can promote HDR in the germline, it should display super-Mendelian inheritance (>50%) of the GFP-marked transgene. All Cas9 sources, which carry the DsRed marker, should have ∼50% inheritance (**Fig. 2a**).

**Fig.2.**
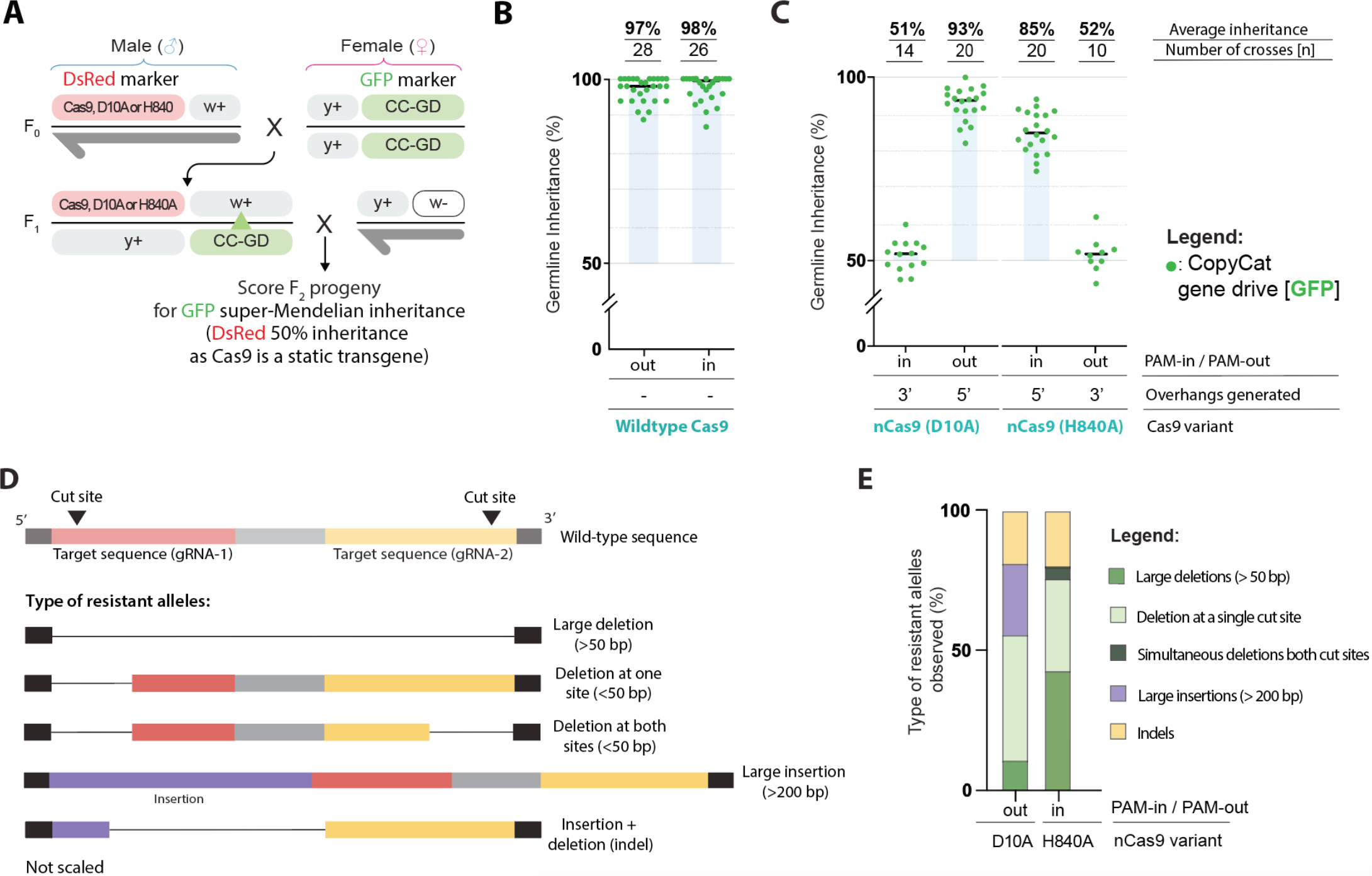
Super-Mendelian inheritance rates produced by nickase Cas9s when 5’ overhangs are generated. **a**. All Cas9 sources (widltype Cas9, nD10A, and nH840A) and the CopyCat elements are inserted in the X chromosome (*yellow [y]* and *white [w]* genes, respectively). F0 males containing the Cas9 were crossed to females containing either Copycat gene drives (CC-GD). F1 females carrying both transgenes were crossed to wildtype males to assess germline allelic conversion (green triangle indicates potential wildtype allele replacement) by scoring the GFP marker in the F2. **b**. Similar biased inheritance rates were observed when widltype Cas9 was combined with both CopyCat elements. **c**. nD10A and nH840A triggered super-Mendelian inheritance rates only when generating 5’ overhangs. **d**. Schematic of observed resistant allele outcomes in the nickase-based gene-drive experiments. **e**. nD10A (n = 20) produced a high frequency of large insertions while nH840A (n = 24) produced bigger deletions between nick sites.

When combining the wildtype Cas9 with both CopyCat elements, we introduce two proximal DNA double-strand breaks by a multiplexing approach (**Fig.1e**), which efficiently biases Mendelian inheritance (Champer et al., 2018). Here, we observed similar super-Mendelian inheritance levels of ∼97% with both PAM-out and PAM-in CopyCat elements (**Fig. 2b**). We did not observe significant differences between CopyCat elements when combined with the widltype Cas9, which suggested that both pairs of gRNA would have similar efficiencies (*p* = 0.6721, see statistics **Supplementary Data 1**). In previous work, we individually validated the *w2*-gRNA in a similar CopyCat arrangement that showed 90% super-Mendelian inheritance (López Del Amo et al., 2020a). The addition here of a second gRNA (*w8* or *w9*) boosted the allelic conversion efficiency from 90% to ∼97% in both cases. These results concur with reports of increased gene-drive performance with an additional gRNA (Champer et al., 2018).

After confirming the activity of the two elements built to test the nickase gene drive, we evaluated the ability of nCas9 to promote super-Mendelian inheritance in either of the four defined scenarios (**Fig. 1e**). First, we crossed our nD10A transgenic line with both CopyCat strains carrying the tandem gRNAs and followed the same experimental cross scheme (**Fig. 2a**). In this case, the nD10A version displayed 93% super-Mendelian inheritance levels when combined with the CC(*w2, w8*) that generated 5’ overhangs. In contrast, the CC(*w2, w9*) (PAM-in) that generated 3’ overhangs was inherited in a Mendelian fashion (∼50%) (**Fig. 2c; Supplementary Data 2**).

Next, we tested the nH840A line following the same cross scheme (**Fig. 2a**). On average, nH840A produced super-Mendelian inheritance rates of ∼85% when combined with the CC(*w2, w9*) (PAM-in) that generated 5’ overhangs. However, we observed 50% inheritance rates when combined with the CC(*w2, w8*) (PAM-out) to generate 3’ overhangs (**Fig. 2c; Supplementary Data 2**). Although we observed super-Mendelian inheritance when using both nD10A and nH840A, nD10A produced biased inheritance only when combined with the CC(*w2, w8*). In contrast, nH840A triggered super-Mendelian inheritance when crossed to the CC(*w2, w9*). We detected gene-drive activity only in these combinations, which generated 5’ overhangs. These results concurred with previous *in vitro* studies where paired nicks only stimulated significant HDR levels when using nD10A to generate 5’ overhangs(Mali et al., 2013; Ran et al., 2013).

Overall, nD10A performed significantly better than nH840A for inheritance bias (*p* < 0.0001, see statistics **Supplementary Data 2**), which could be due to different cleavage rates between the nickases. The *white* gene targeted for conversion in our system is tightly linked with the *yellow* gene where our nCas9 sources are inserted, which allowed us to evaluate the cutting rates via the F_2_ progeny. In fact, nD10A displayed 95% cutting efficiency, which is higher than nH840A that showed 91% cutting rates (*p* = 0.0683; see statistics **Supplementary Data 1**). Altogether, we have demonstrated the first example of a gene-drive system driven by a nCas9, which simultaneously nicks both complementary strands to induce efficient allelic conversion that is mediated by HDR *in vivo*. Furthermore, our data indicates that super-Mendelian inheritance through nickase gene drives can only be achieved by generating 5’ overhangs.

DNA double-strand breaks introduced by gene drives can produce resistant alleles at the target site when the HDR-mediated allelic conversion process is inaccurate. Resistant alleles generated by wildtype Cas9 have been characterized by us and other groups (Champer et al., 2018; Gantz and Bier, 2015; Hammond et al., 2017; López Del Amo et al., 2020b). However, the types of resistant alleles generated by paired nicks are uncharacterized in a gene-drive context. Thus, we identified and evaluated the resistant alleles generated by our nickase gene-drive systems. Since the CopyCat target is the *white* gene located in the X chromosome, and males have only one X chromosome, GFP-negative F_2_ males that present the *white* eye phenotype indicate *white* gene disruption and unsuccessful allelic conversion, indicating individuals that carry resistant alleles. Thus, we collected non-converted F_2_ males from the nickase conditions that displayed gene-drive activity, nD10A PAM-out and nH840 PAM-in (**Fig. 2c**). We extracted the DNA from these animals and characterized them by Sanger sequencing. We observed five different classes of resistant alleles for both PAM-out and PAM-in conditions: i) large deletions spanning both cut sites (>50 base pairs [bp]), ii) deletions (<50 bp) occurring at one cut site with the other cut site intact, iii) simultaneous deletions (<50 bp) at both target sites with partial sequence retention between cut sites, iv) large insertions (>200 bp) and v) small insertions combined with small deletions in the same individuals at the same cut site (i.e., indels) (**Fig. 2d; Supplementary Fig.1**).

We detected that nH840A combined with PAM-in gRNAs comprised 42% of the total sequenced flies with large deletions spanning 50 to 90 nucleotides (**Fig. 2e; Supplementary Fig. 1a**). This is a significantly higher frequency of large deletions spanning both nick sites compared to the PAM-out (5%) (**Fig. 2e;** *p* = 0.0389, see statistics **Supplementary Data 3**). We also observed large insertions (>200 bp) when nD10A was combined with paired gRNAs in a PAM-out orientation (**Supplementary Fig.1d**). We found that 25% of the sequenced flies within this condition had large insertions, which was significantly higher than nH840A combined with PAM-in gRNAs, where we did not detect any large insertion event (**Fig. 2e**; *p* = 0.0143, see statistics **Supplementary Data 3**). Interestingly, the large insertions produced by the PAM-out gRNAs represented partial HDR occurrences containing portions of the engineered gene-drive allele. We detected either part of the U6 promoter downstream of the left homology arm or the synthetic 3xP3 promoter with an incomplete piece of the GFP marker downstream of the right homology arm of the gene-drive element (**Supplementary Fig. 1d**). This aligned with previous studies that used a multiplexing gene drive with wildtype Cas9 and gRNAs in a PAM-out orientation that induced the formation of insertions resulting from partial HDR events (Champer et al., 2018).

We did not observe significant differences between the nickase genotypes in the other categories described above, including single cuts at only one target site, simultaneous mutations at both cut sites or indels (**Fig. 2d-e**; see statistics **Supplementary Data 3**). We sequenced the resistant alleles generated by the widltype Cas9 and confirmed that the gRNAs in this work are active, as we found mutations at all target sites within the genotypes that produced super-Mendelian inheritance with the widltype Cas9 and nCas9s (**Supplementary Fig.1**). Overall, we have shown different repair outcomes when using distinct nickases, which can inform future nickase-based gene-drive strategies.

## DISCUSSION

In this work, we describe a novel gene-drive system based on nickase versions of Cas9 that promotes super-Mendelian inheritance in *Drosophila*, and increases the number of feasible design options for gene drives aimed at population engineering. We showed that both nD10A and nH840A produced efficient HDR in the germline, but only when the two nicks in the DNA generated 5’ overhangs. We characterized events that failed to convert the wildtype allele to the gene drive by Sanger sequencing, which indicated that nH840A combined with PAM-in gRNAs produced higher rates of large deletions compared to the nD10A and PAM-out arrangement. While the PAM-out condition triggered large insertions, we did not observe such alterations when we analyzed resistant alleles produced by the nH840A, which suggested that the modes of DNA repair that are triggered are nickase dependent.

In our experiments, nD10A produced higher super-Mendelian rates than nH840A, which we attributed to differences in cleavage activities between the nickases. In fact, nH840A has been shown to present less cleavage activity *in vitro* as it produced lower indel rates when disrupting the *EMX-S1* gene(Gopalappa et al., 2018). Additionally, the distinct time-windows of cleavage between the pairs of gRNAs, which need to cut simultaneously, may impact HDR efficiencies as both nickases showed super-Mendelian inheritance with different paired gRNAs. While previous studies did not report meaningful HDR rates with nH840A (Bothmer et al., 2017; Mali et al., 2013; Ran et al., 2013), we showed efficient HDR rates achieved by gRNAs in a PAM-in configuration when generating 5’ overhangs using nH840A for the first time. Thus, nH840A could be a viable option for future nickase-based designs to promote HDR.

While the focus of this work was to obtain a proof-of-concept for a Cas9-nickase gene drive, we observed slightly higher super-Mendelian rates in the wildtype Cas9 over the nickases. We believe that this is because a nickase requires the coordinated action of both gRNAs cutting simultaneously to induce efficient HDR. Conversely, when the wildtype Cas9 is used, a single cut from either of the paired gRNAs can induce HDR to produce double-stranded DNA breaks. Furthermore, while failure of the first gRNA would result in a small indel, the second gRNA can still cut and trigger a second round of potential conversion as previously shown (Champer et al., 2018), and which would explain the higher inheritance rates in the wildtype Cas9 scenario. Thus, it is imperative to thoroughly test the paired gRNAs to maximize coordinated action in future nickase gene-drive systems.

With regard to resistant allele formation, we have shown that our nH840A transgene combined with paired gRNAs in a PAM-in configuration frequently generated large deletions between the spaced nicks, which we did not detect when using gRNAs in a PAM-out configuration with nD10A. We can harness this effect to boost gene-drive efficiency when specifically targeting essential genes. For example, the gene-drive element can carry a DNA rescue sequence to replace a wildtype allele while restoring functionality of vital genes to ensure animal viability and gene-drive spread. If resistant alleles occur from unsuccessful allelic conversion, these mutations should produce non-viable animals that evade propagation, though small mutations in essential genes can still produce some viable escapees (Terradas et al., 2021). Therefore, large deletions induced by a gene-drive system using nH840A in a PAM-in configuration should help remove surviving escapees in a population carrying small indels.

We questioned whether a nickase gene-drive system could reduce the formation of resistant alleles, as DNA nicks are usually repaired efficiently. We detected resistant alleles caused by single DNA nicks, which could be from non-repaired single nicks that were converted to double-strand breaks that were subsequently fixed by non-homologous end joining (Kuzminov, 2001). Importantly, mutations produced at a single target site by DNA nicks could limit further gene-drive propagation, as single nicks are poor HDR inducers (Vriend et al., 2016) and a single mutation at one target site would be enough to avoid gene-drive spread. However, our proposed nickase system is amenable to further optimization, especially as one major contributing factor to this might be due to fixing the induced DNA paired nicks to ∼50 nucleotides apart, though in fact, efficient HDR has been observed with offset distances ranging from 20 to 100 base pairs (Vriend et al., 2016). Thus, we envision further adapting our strategy in the future to generate distinct offset distances to improve the HDR efficiency of our nickase-based system.

Future nickase-gene-drive approaches could explore additional optimizations to increase specificity and reduce off-target effects, which can accumulate in a population and must be considered. As two independent cleavage events need to be coordinated for the desired modification, paired nicks were shown in previous work to improve specificity while reducing off-target effects when disrupting the *EMX1* gene in human cell culture (Ran et al., 2013). Recently, the off-target effects of gene drives were predicted using validated algorithms and posterior *in vivo*-targeted deep sequencing with *Anopheles* mosquitoes in laboratory cage studies (Garrood et al., 2021). The off-targets effects were almost undetectable if using promoters that restricted Cas9 expression to the germline. Indeed, a nickase gene-drive system could be tested in *Anopheles* and species with much larger genomes, such as *Aedes* or *Culex* mosquitoes (Main et al., 2021; Severson et al., 2004), to study the pervasiveness of off-target effects across genome sizes and in the wild. Furthermore, gene drives can bias Mendelian inheritance in mice (Grunwald et al., 2019; Weitzel et al.). In the future, a nickase-based gene-drive system could also be applied to mice or to reduce off-target effects when editing mammalian embryos (Aryal et al., 2018).

Altogether, our proof-of-principle study provides a first step towards the development of next-generation nickase-based gene drives that advances their potential future applications. We envision that our work will spur the use of nickase-based gene-drive systems for improved population control while encouraging its implementation in a broader range of organisms.

## ACKNOWLEDGMENTS

We thank Kaycie Butler and JingXin Liang for comments and edits on the manuscript. The research reported in this manuscript was supported by the University of California, San Diego, Department of Biological Sciences, the Office of the Director of the National Institutes of Health under award number DP5OD023098 (to V.M.G), and the National Institute of Allergy and Infectious Diseases under the award number R01AI162911 (to V.M.G). V.L.D.A and V.M.G are authors on a patent filed by the University of California, San Diego (Provisional Application Serial No. 63/254,753) that is related to the nickase gene-drive system described in this work.

## AUTHOR CONTRIBUTIONS

V.L.D.A and V.M.G conceived the project. V.L.D.A designed and obtained the nickase gene-drive constructs in *Drosophila*. V.L.D.A and S.S.J performed the experiments. V.L.D.A, V.M.G and S.S.J performed the figure visualizations. V.L.D.A wrote the manuscript, which was edited by all the authors.

## MATERIAL AND METHODS

### EXPERIMENTAL MODEL AND SUBJECT DETAILS

#### Fly rearing and crosses

All flies were kept on standard food with a 12/12 hours day/night cycle. Fly stocks are kept at 18°C, and all experimental crosses were performed at 25°C. All flies were anesthetized during our experiments using CO_2_. F_0_ crosses from gene-drive experiments were made in pools of 3-6 virgin females crossed to 3-6 males. F_1_ experiments were always made in single pairs to track editing events happening singularly in the germline. The F_2_ progeny was scored as male or female and sorted for a fluorescent marker (DsRed and/or GFP) using a Leica M165 F2 Stereomicroscope with fluorescence as an indicator of transgene inheritance rates. All experiments were performed in a high-security ACL2 (Arthropod Containment Level 2) facility built for gene drive purposes in the Division of Biological Sciences at the University of California, San Diego. Crosses were made in polypropylene vials (Genesee Scientific Cat. #32-120), and all flies were frozen for 48 hours before being removed from the facility, autoclaved, and discarded as biohazardous waste.

### METHOD DETAILS

#### Plasmid construction

DNA constructs were built using NEBuilder HiFi DNA Assembly Master Mix (New England BioLabs Cat. #E2621) and transformed into NEB 10-beta electrocompetent *E*.*coli* (New England BioLabs Cat. #3020). DNA was extracted using a Qiagen Plasmid Midi Kit (Qiagen Cat. #12143) and sequenced by Sanger sequencing at Genewiz. Primers used for cloning can be found in the Key Resources Table below. Final sequences of all constructs will be available at NCBI ahead of publication.

#### Transgenic line generation and genotyping

We outsourced embryo injections to Rainbow Transgenic Flies, Inc. All DNA constructs were injected into our lab’s isogenized Oregon-R (Or-R) strain to ensure consistent genetic background throughout experiments. Plasmid templates were co-injected with a Cas9-expressing plasmid (pBSHsp70-Cas9 was a gift from Melissa Harrison & Kate O’Connor-Giles & Jill Wildonger [Addgene plasmid #46294; http://n2t.net/addgene:46294; RRID: Addgene_46294]). We received the injected generation 0 (G_0_) animals, then we intercrossed the hatched adults in small pools (3-5 males x 3-5 females), and screened the G_1_ flies for a fluorescent marker (DsRed for Cas9 versions and GFP for gene-drive elements, both fluorescences in the eye), which was indicative of transgene insertion. Lastly, we established homozygous lines from single transformants by crossing to Or-R. As the Cas9 transgene is inserted into the *yellow* gene disrupting it, homozygous flies for the Cas9 versions can be identified once flies display a yellow body color. Similarly for the Copycat constructs that are integrated into the *white* gene, homozygous flies for the CopyCat elements display a white eye phenotype. Stocks were sequenced by PCR and Sanger sequencing to confirm proper transgene insertion.

#### DNA extraction from single flies

To sequence resistant alleles, we extracted genomic DNA from individual males following the method described by Gloor GB and colleagues (Gloor et al., 1993). In brief, we used 50ul of the extraction buffer to squish single flies in a PCR tube. Next, we placed them into the PCR machine (Proflex PCR system, *Applied Biosystems*) for 1 hour at 37°Cfollowed by 5 minutes at 95°C to inactivate the proteinase K. Then, we added to each DNA sample 200uL of water to obtain a total of 250uL per sample. Lastly, we used 1-5uL in a 25uL PCR reaction covering the gRNA cut sites in the *white* gene for Sanger sequencing analysis.

#### Sanger sequencing of individuals carrying resistance alleles

We amplified a DNA region covering the gRNA cut sites using the v1564 and v1565 oligos (see oligos list below). The obtained amplicon was then sequenced by Sanger sequencing to determine the quality of the resistant alleles using the v478 oligo. When we obtained lower-quality traces, we performed a second Sanger sequencing reaction from the other side of the amplicon to confirm the quality of the mutation with either the v659 or v1571 primers. Primers used for resistance allele sequencing can be found in the Key Resources Table.

#### Microscopy

Adult flies were anesthetized using CO2 to select individuals for crossing experiments. Their phenotypes were analyzed using a Leica M165 FC Stereo microscope to properly prepare the experimental crosses. Inheritance analysis of the transgenes marked with fluorescence was evaluated using the same microscope. DsRed marker implies presence of the Cas9 cassettes while GFP fluorescence indicates presence of the CopyCat transgenes.

### QUANTIFICATION AND STATISTICAL ANALYSIS

#### Statistical analysis

We used GraphPad Prism 9 and Adobe Illustrator to generate all the graphs. Statistical analyses were performed using GraphPad Prism 9. In **Fig. 2b**, we applied an unpaired *t test* to compare inheritance rates when using the wildtype Cas9 (**Supplementary Table 1**). Additionally, One-Way Anova and Tukey’s multiple comparison test to evaluate differences between super-Mendelian rates in our nickase gene-drive experiments in **Fig. 2c** (**Supplementary Table 2**). For evaluating the differences in proportions for resistant allele events in **Fig. 2e**, we used Fisher’s exact test (**Supplementary Table 3**).

## KEY RESOURCES TABLE

**Supplementary Table 1.**
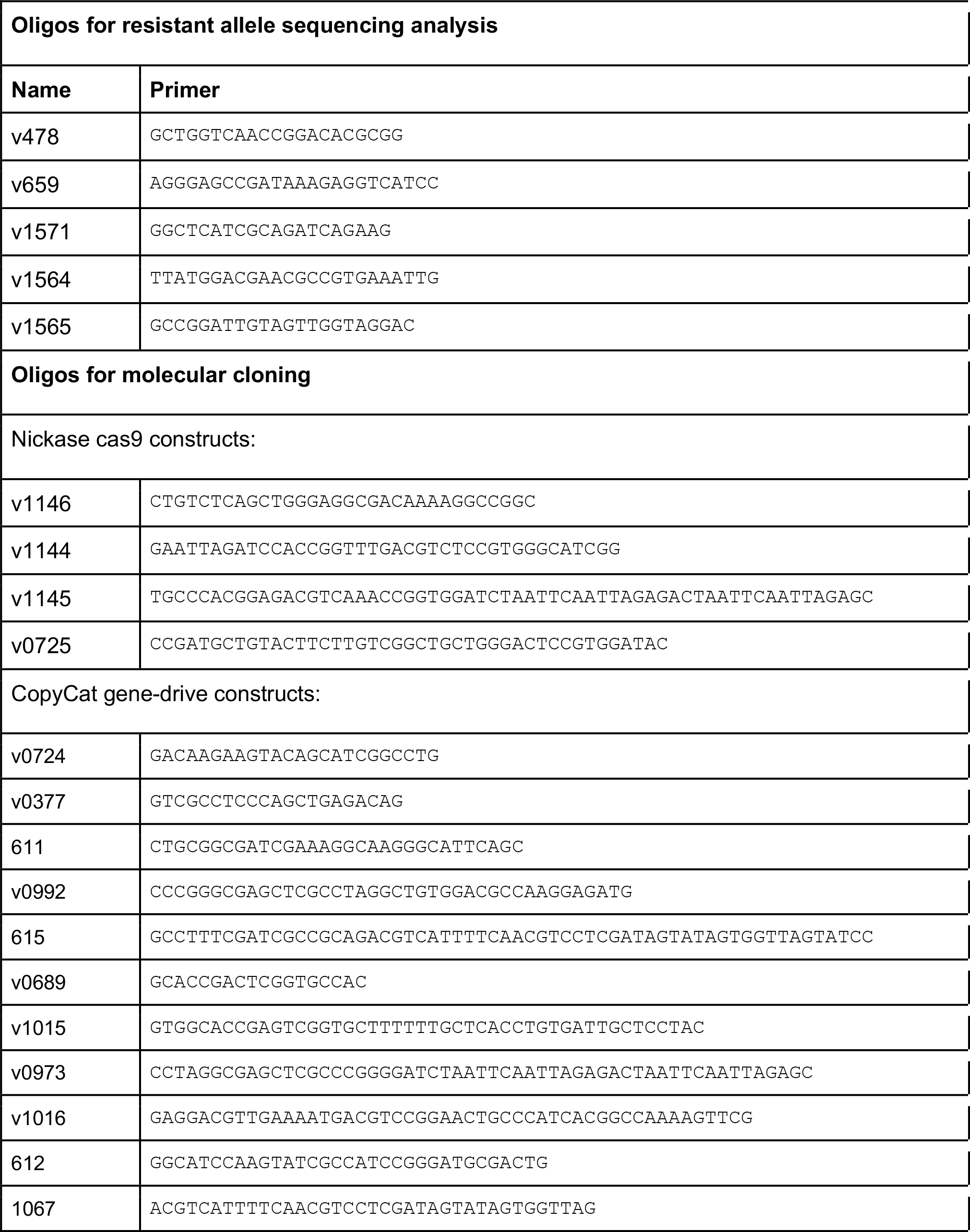
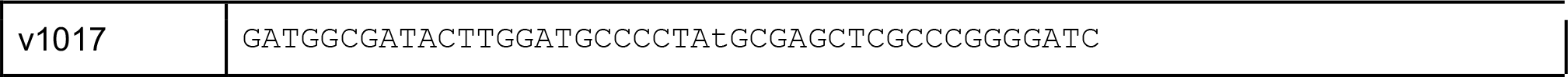
List of primers used in this study.

## SUPPLEMENTAL TABLES

**Data S1.xlsx, Related to Figure 2**. Phenotypical scoring of the two CopyCat gene-drives elements combined with wildtype cas9.

**Data S2.xlsx, Related to Figure 2**. Phenotypical scoring of the two CopyCat gene drives combined with either nickase Cas9 version.

**Data S3.xlsx, Related to Figure 2 and Supplementary Figure 1**. Comparison of deletion and insertion events between both nickase versions for resistant allele formation.

