## Supplementary Figure 1 for "A nickase Cas9 gene-drive system promotes super-Mendelian inheritance in *Drosophila*"

### (A) Large deletions (>50nt)

#### nD10A PAM-out

```
WT  :  CGATCGCCGCAG   GGCATCCAAGTATCGCCATCCGGGATGCGACTGCTCAATGGCCAAC   CTGTGGACGCCAA

       CGATC·······   ··············································   CTGTGGACGCCAA   (-53bp)
       CGATCGCCGCAG   GGC···········································   ··········CAA   (-53bp)
```

#### nH840A PAM-in

```
WT  :  TTGGCCGTGATGGGCAGTTCCGG   TGCCGGAAAGACGACCCTGCTGAATGCCCTTGCCTTTCGATCGCCGCAG   GGCATCCAAGTATCGCCATCCGGGATG

       TTGGCCGTGATGGGCAGTTCCGG   ·················································   ··CATCCAAGTATCGCCATCCGGGATG   (-51bp)
       TTGGCCGTGATGGGCAGTTCC··   ·················································   ···ATCCAAGTATCGCCATCCGGGATG   (-54bp)
       TTGGCCGTGATGGGCAGTTCC··   ·················································   ·······AAGTATCGCCATCCGGGATG   (-58bp)
       TTGGCCGTGATGGGCAGTT····   ·················································   ·····CCAAGTATCGCCATCCGGGATG   (-59bp)
       TTGGCCGTGATGGGCAGT·····   ·················································   ···········ATCGCCATCCGGGATG   (-65bp) (X4)
       TTG····················   ·················································   ···························   (-96bp)
       TTGG···················   ·············ACCCTGCTGAATGCCCTTGCCTTTCGATCGCCGCAG   GG······················ATG   (-54bp)
```

#### Cas9 PAM-out

```
WT  :  CGATCGCCGCAG   GGCATCCAAGTATCGCCATCCGGGATGCGACTGCTCAATGGCCAAC   CTGTGGACGCCAAGG

       CGATCGCCGCAG   ··············································   ···············   (-61bp)
```

#### Cas9 PAM-in

```
WT  :  TTGGCCGTGATGGGCAGTTCCGG      TGCCGGAAAGACGACCCTGCTGAATGCCCTTGCCTTTCGATCGCCGCAG    GGCATCCAAGTATCGCCATCCGGGATG

       TTGGCCGTGATGGGCAGTTCCGG      .................................................    ..CATCCAAGTATCGCCATCCGGGATG   (-51bp)
       TTGGC..................      .................................................    ...ATCCAAGTATCGCCATCCGGGATG   (-70bp) (X3)
```

### (B) Cuts at a single target site

#### nD10A PAM-out

```
WT  :  CGATCGCCGCAG      GGCATCCAAGTATCGCCATCCGGGATGCGACTGCTCAATGGCCAAC      CTGTGGACGCCAA

       CGATCGCCGCAG      GGCATCCAAGTATCGCCATCCGGGATGCGACTGCTCAATGGCCAAC      ........GCCAA       (-8bp)
       CGATCGCCGCAG      .........GTATCGCCATCCGGGATGCGACTGCTCAATGGCCAAC      CTGTGGACGCCAA       (-9bp)
       CGATCGCCGCAG      G.......................ATGCGACTGCTCAATGGCCAAC      CTGTGGACGCCAA       (-23bp)
       CGATCGCCGCAG      MP                          ...................     ...TGGACGCCAA       (-22bp)
       CGATCGCCGCA       MP                          ...................     ...TGGACGCCAA       (-22bp)
       CGATCGCCGCAG      MP                      .......................     CTGTGGACGCCAA       (-23bp)
       CGATCGCCGCAG      MP                         ....................     ..GTGGACGCCAA       (-23bp)
       CGATCGCCGCA       MP                      .......................     ...TGGACGCCAA       (-26bp)
       CGATCGCCGCA       MP                  ...........................     .TGTGGACGCCAA       (-27bp)
```

#### nH840A PAM-in

```
WT  :  TTGGCCGTGATGGGCAGTTCCGG      TGCCGGAAAGACGACCCTGCTGAATGCCCTTGCCTTTCGATCGCCGCAG    GGCATCCAAGTATCGCCATCCGGGATG

       TTGGCCGTGATGGGCAGTTCCGG      TGCCGGAAAGACGACCCTGCTGAATGCCCTTGCCTTTCGATCGCCGCAG    G. MP                         (-1bp)
       TTGGCCGTGATGGGCAGTTCCGG      .....GAAAGACGACCCTGCTGAATGCCCTTGCCTTTCGATCGCCG       MP                            (-5bp)
       TTGGCCGTGATGGGCAGTTCCGG      MP                                              .    ....TCCAAGTATCGCCATCCGGGATG   (-5bp)
       TTGGCCGTGATGGGCAGTTCCGG      TGCCGGAAAGACGACCCTGCTGAATGCCCTTGCCT MP     .......    .....CCAAGTATCGCCATCCGGGATG   (-12bp)
       TTGGCCGTGATGGGCAGTT          MP                          ...................AG    MP                            (-18bp) (x2)
       TTGGCCG.................     .....G                                               MP                            (-21bp)
       TTGGCCGTGATGGGCAGTTC         MP                        .....................     ...ATCCAAGTATCGCCATCCGGGATG   (-25bp)
```

#### Cas9 PAM-out

```
WT  :  CGATCGCCGCAG      GGCATCCAAGTATCGCCATCCGGGATGCGACTGCTCAATGGCCAAC      CTGTGGACGCCAAGG

       CGATCGCCGCAG      G·CATCCAAGTATCGCCATCCGGGATGCGACTGCTCAATGGCCAAC      CTGTGGACGCCAAGG      (-1bp)
       CGATCGCCGCAG      ··CATCCAAGTATCGCCATCCGGGATGCGACTGCTCAATGGCCAAC      MP                   (-2bp)  (X2)
```

### (C) Simultaneous cuts at both target sites

#### nH840A PAM-in

```
WT  :  TTGGCCGTGATGGGCAGTTCCGG      TGCCGGAAAGACGACCCTGCTGAATGCCCTTGCCTTTCGATCGCCGCAG      GGCATCCAAGTATCGCCATCCGGGATG

       TTGGCCGTGATGGGCAGTTCCGG      ·············ACCCTGCTGAATGCCCTTGCCTTTCGATCGCCGCA·      ····TCCAAGTATCGCCATCCGGGATG   (-18bp)
```

#### Cas9 PAM-out

```
WT  :  CGATCGCCGCAG      GGCATCCAAGTATCGCCATCCGGGATGCGACTGCTCAATGGCCAAC      CTGTGGACGCCAAGG

       CGATCGCCGCAG      G·CATCCAAGTATCGCCATCCGGGATGCGACTGCTCAATGGCC···      ·TGTGGACGCCAAGG      (-5bp)
       CGATCGCCGCAG      G···TCCAAGTATCGCCATCCGGGATGCGACTGCTCAATGGCCAA·      ·TGTGGACGCCAAGG      (-5bp)
       CGATCGCC····      ·················ATCCGGGATGCGACTGCTCAATGGCCAA·      ·TGTGGACGCCAAGG      (-23bp)
       CGATCGCCGCAG      G····························ACTGCTCAATGGCCAAC      ·TGTGGACGCCAAGG      (-29bp)
       CGATCGCCGCAG      GG·····················GATGCGACTGCTCAATGGCCAA·      ·············GG      (-36bp)
```

#### Cas9 PAM-in

```
WT  :  TTGGCCGTGATGGGCAGTTCCGG      TGCCGGAAAGACGACCCTGCTGAATGCCCTTGCCTTTCGATCGCCGCAG      GGCATCCAAGTATCGCCATCCGGGATG

       TTGGCCGTGATGGGCAGTTCCG·      TGCCGGAAAGACGACCCTGCTGAATGCCCTTGCCTTTCGATCGCCGCAG      G·CATCCAAGTATCGCCATCCGGGATG   (-2bp)  (x6)
       TTGGCCGTGATGGGCAGTTCCGG      ······AAAGACGACCCTGCTGAATGCCCTTGCCTTTCGATCGCCGCAG      G·CATCCAAGTATCGCCATCCGGGATG   (-7bp)  (X4)
       TTGGCCGTGATGGGCAGT·····      ·GCCGGAAGACGACCCTGCTGAATGCCCTTGCCTTTCGATCGCCGCAG       G·CATCCAAGTATCGCCATCCGGGATG   (-7bp)
       TTGGCCGTGATGGGCAGTTCCGG      ······AAAGACGACCCTGCTGAATGCCCTTGCCTTTCGATCGCCGCAG      G··ATCCAAGTATCGCCATCCGGGATG   (-8bp)  (X2)
```

### (D) Large insertions (>200bp)

### nD10A PAM-out

**WT : CGATCGCCGCAG GGCATCCAAGTATCGCCATCCGGGATGCGACTGCTCAATGGCCAAC CTGTGGACGCCAAGG**

CGATCGCCGCAG ACGTCATTTTCAACGCCATCCATGGTATGAGGGCTAATATCCCCGCCTGTGACGCGGGAGAAAAGGGGGGAAATG
CCCGCCTGGGAGCATCAGGAATTCTTTTTTATNTACTTTANNNNNGTTATATAAACAATTTTTGTTTTAATTGAATCTAATTTGCCATTGCTTTTAG
GAATCTCANGCATCCANCAAGCGTTTGTCCGCCGAATCGCCCNTCANTGAANAAGATCCTGTGGCG (+238bp) HDR (U6)

CGATCGCCGCAG ACGTCATTTTCAACGTCCTCGATAGTATAGTGGTTAGTATCCCCGCCTGTCACGCGGGAGACCGGGGTTCAATTC
CCGTCGGGGAGAATCTGTGATTCTTTTTTTTTTTCTTTTACTTTGTTATATAAACAATTTTTGTTTTAATTGAATCTAATTTGCCATTGCTTTTAGG
AATCTCAGGCATCCAGCAAGCGTTTGTCCGCCGAATCGCCCATCAGTGAAGAAGATCCTGTGGCGGCTACGAAAATCTCCCCGGCCATGTCGGCCTC
CACCTCCAGCGAAAAACCCATCAGCGAGCTGGCCACCTCTGTGCTGACCCACCGCTTTCCAGACTCCACCTCCTCACCCGGCGAACATGGCCTTGGA
CGAATGCAGTTGTCGATCCGCTACAGCGCCCAGCGTCAAAAACTAGACGTGACCATACACAAAATCCAGAAGATACCACTTCGCGATCCCAGCAA (+462bp) HDR (U6)

CGATCGCCGCGA ACGTCATTTTCAACGTCCTCGATAGTATAGTGGTTAGTATCCCCGCCTGTCACGCGGGAGACCGGGGTTCAAT
TCCCCGTCGGGGAGAATCTGTGATTCTTTTTTTTTTTCTTTTACTTTGTTATATAAACAATTTTTGTTTTAATTGAATCTAATTTGCCATTGCTTTT
AGGAATCTCAGGCATCCAGCAAGCGTTTGTCCGCCGAATCGCCCATCAGTGAAGAAGATCCTGTGGCGGCTACGAAAATCTCCCCGGCCATGTCGGC
CTCCACCTCCAGCGAAAAACCCATCAGCGAGCTGGCCACCTCTGTGCTGACCCACCGCTTTCCAGACTCCACCTCCTCACCCGGCGAACATGGCCTT
GGACGAATGCAGTTGTCGATCCGCTACAGCGCCCAGCGTCAAAAACTAGACGTGACCATACACAAAATCCAGAAGATACCACTTCGCGATCCCAGCA
ATATCCCCGATCCGTATGTTAAGCTGTATCTGTTGCCTGGACGCACCAAGGAGTCGAAACGCAAGACGAGCGTGATCAAGGACAACTGCAACCCCGT
CTACGATGCATCCTTTGAGTACCTGATTTCCATTGCCGAACTCAGGCAGACGGAACTGGAGGTGACGGTGTGCACCCAAAAGGGATTCCTATCCGGC
GGTAGTCCCATCATTGGCATGGTAGGTACCCGAAAGCAACCCCTTAGTTACAGACNCAGCGCGTACGTCCTTCGCATCCTTATGATTCCCAAGTACA
TATNTGCANANTACAGTATATATAGGAAAGANATCCNGNGAACTTCGNCGATACTTGGNTGCCCTGGTTTANAGCTA (+828bp) HDR (U6)

CGATCGCCGCAG MP · AGGCAATGGTCCATGGGTCAATGGGCGAAGGCCTTCAAGTCGATGGGGGTGA
CCNGGGTGGCCCCCTCNAACTTCCCCTCCGCCCGGGTCATANAATTGCCGTCGTCCTTGAAGAAGATGGTGCGCTCCTGGACGTAACCTTCGGGCAT
GGCGGACTTGAAAAAGTCGTGCTGCTTCATGTGGTCGGGGTAGCGGCTGAAGCACTGCACGCCGTANGNCAGGGTGGTCACCAAGGTGGGCCANGGC
ACGGGCAGCTTGCCGGTGGTGCANANGAACTTCANGGTCAGCTTGCCGTANGTGGCATCGCCCTCNCCCTCGCCGGACACGCTGAACTTGTGGGCGT
TTACNTCGCCGTCCANNTCGACCANGATGGGCACCACCCCGGTGAACANCTNCTCGCCCTTGNTCACCATGGTGGCGANCGGTGGATCCCGGGCCCG
CNGGTACCGTCNACTCTAGCGGNACCCCCNNNTGNNTNAGCTTGTTCAGCTGCGCTTGNTTNTTTGCNTAGCTTTCGCTTAGCGGANGTGNTCACTT
TGCTTGGTTTGAATTGGAATGTCGCTCCNTAGACNAAGCGCCTCTATTTATACTNCGGCNGTCGANGGTTCGAAATCGATNAGCTTGGANCTAATGA
ATAGCTCTAATGANT (-1bp) (+649) HDR (3xP3)

CGATCGCCGCAG GGCATCCAAGTATCGCCATCCGGGATGCGATGGCTCAATGGCCAA GGGCCAGGGCACGGGCAGCTT
GTTGCCGGTGGTGCAGATGAACTTCAGGGTCAGCTTGCCGTAGGTGGCATCGCCCTCGCCCTCGCCGGACACGCTGAACTTGTGGCCGTTTACGTC
GCCGTCCAGCTCGACCAGGATGGGCACCACCCCGGTGAACAGCTCCTCGCCCTTGCTCACCATGGTGGCGACCGGTGGATCCCGGGCCCGCGGTAC
CGTCGACTCTAGCGGTACCCCGATTGTTTAGCTTGTTCAGCTGCGCTTGTTTATTTGCTTAGCTTTCGCTTAGCGACGTGTTCACTTTGCTTGTTT
GAATTGAATTGTCGCTCCGTAGACGAAGCGCCTCTATTTATACTCCGGCGGTCGAGGGTTCGAAATCGATAAGCTTGGATCCTAATTGAATT
AGCTCTAATTGAATTAGTCTCTAATTGAATTAGATCCCCGGGCGAGCTCGCCTANG · CTGTGGACGCCAAGG (-1bp) (+452bp) HDR (3xP3)

**Cas9 PAM-out**

```
WT   :  CGATCGCCGCAG           GGCATCCAAGTATCGCCATCCGGGATGCGACTGCTCAATGGCCAAC       CTGTGGACGCCAAGG

        CGATCGCCGCAG           G·CATCCAAGTATCGCCATCCGGGATGCGACTGCTCAATGG
        ACTTGTGGCCGTTTACGTCGCCGTCCAGCTCGACCAGGATGGGCACCACCCCGGTGAACAGCTCCTCGCCCTTGCTCACCATGGTGGCGAC
        CGGTGGATCCCGGGCCCGCGGTACCGTCGACTCTAGCGGTACCCCGATTGTTTAGCTTGTTCAGCTGCGCTTGTTTATTTGCTTAGCTTTC
        GCTTAGCGACGTGTTCACTTTGCTTGTTTGAATTGAATTGTCGCTCCGTAGACGAAGCGCCTCTATTTATACTCCGGCGGTCGAGGGTTCG
        AAATCGATAAGCTTGGATCCTAATTGAATTAGCTCTAATTGAATTAGTCTCTAATTGAATTAGATCCCCGGGCGAGCTCGCCTAGG
                                                                                 ·····           CTGTGGACGCCAAGG    (-6bp)(+359) HDR (3xP3)

        CGATCGCCGCAG           TTTGACCATAGTGTTTCAATTCTACATTAATTTTACAGAGTAGAATGAAACGCCACCTACTCAGCCAA
        GAGGCGAAAAGGTTAGCTCGCCAAGCAGAGAGGGCGCCAGTGCTCACTACTTTTTATAATTCTCAACTTCTTTTTCCAGACTCAGTTCGTA
        TATATAGACCTATTTTCAATTTAACGTAACATCAAATTTTCTGTCAATAAAGCATATTTATTTATATTTATTTTACAGGAAAGAATTCCTT
        TTAAAGTGTATTTTAACCTATAATGAAAAACGATTAAAAAAAAATACATAAAATAATTCGAAAATTTTTGAATAGCCCAGGTTGATAAAAA
        TTCATTTCATACGTTTTATAACTTATGCCCNTAAGTATTT          CGACTGCTCAATGGCCAAC      CTGTGGACGCCAAGG    (+382bp) HDR (U6)

        CGATCGCCGCAG           CAACTGCAACCCCGTNTACGATGCATCCTTTGAGTACNTGATTTCCATTGCCGAACTCAGGCAGACGG
        AACTGGAGGTGACGGTGTGCACCCAAAAGGGATTCCTATCCGGCGGTAGTCCCATCATTGGCATGGTAGGTACCCGAAAGCAACCCCTTAG
        TTACAGACACAGCGCGTACGTCCTTCGCATCCTTATGATTCCCAAGTACATATTCTGCAAGAGTACAGTATATATAGGAAAGATATCCGGG
        TGAACTTCGGCGATACTTGGATGCCCTGGTTTTAGAGCTAGAAATAGCAAGTTAAAATAAGGCTAGTCCGTTATCAACTTGAAAAAGTGGC
        ACCGAGTCGGTGCTTTTTTGCTCACCTGTGATTGCTCCTACTCAAATACAAAAACATCAAATTTTCTGTCAATAAAGCATATTTATTTATA
        TTTATTTTACAGGAAAGAATTCCTTTTAAAGTGTATTTTAACCTATAATGAAAAACGATTAAAAAAAAATACATAAAATAATTCGAAAATTT
        TTGAATAGCCCAGGTTGATAAAAATTCATTTCATACGTTTTATAACTTATGCCCCTAAGTATTTTTTGACCATAGTGTTTCAATTCTACAT
        TAATTTTACAGAGTAGAATGAAACGCCACCTACTCAGCCAAGAGGCGAAAAGGTTAGCTCGCCAAGCAGAGAGGGCGCCAGTGCTCACTAC
        TTTTTATAATTCTCAACTTCTTTTTCCAGACTCAGTTCGTATATATAGACCTATTTTCAATTTAACGT
                                                          CGACTGCTCAATGGCCAAC      CTGTGGACGCCAAGG     (+773bp) HDR (U6)
```

### (E) Indels

#### nD10A PAM-out

```
WT  :  CGATCGCCGCAG        GGCATCCAAGTATCGCCATCCGGGATGCGACTGCTCAATGGCCAAC      CTGTGGACGCCAAGG

       CGATCGCCGCAG        G·············································      ···TGGACGCCAAGG          (-49bp) (+3bp)
                            AGA
       CGATCGCCGCA·        ··············································      ··GTGGACGCCAAGG          (-49bp) (+3bp)
                 ATC
       CGATCGCCGCAG        GGCATCCAAGTATCGCCATCCGGGATGCGATTGTTGT·TGG···AC      ·TGTGGACGCCAAGG          (-5bp)  (+4bp) (S 4bp)
                                                      GAAC
       CGATCGCCG··    ·    ·GCAT···········TTCCGGGATGCGATGGCTCAATGGCCAAC      CTGTGGACGCCAAGG          (-15bp) (+3bp) (S 1bp)
                                                     CCC
```

#### nH840A-PAM-in

```
WT  :  TTGGCCGTGATGGGCAGTTCCGG      TGCCGGAAAGACGACCCTGCTGAATGCCCTTGCCTTTCGATCGCCGCAG   GGCATCCAAGTATCGCCATCCGGGATG

       TTGGCCGTGATGGGCAGTTCCGG      MP                                             A···   ·······AAGTATCGCCATCCGGGATG   (-10bp) (+1bp)
       TTGGCCGTGATGGGCAGTTCCGG      TTGCCGGAAAGACGACCCTGCTGAATGCCCTTGCCTTTCGATCGCCGCAG   G·········· MP                (-10bp) (+1bp)
       TTGGCCGT···············      ·  ·················································   GGCATCCAAGTATCGCCATCCGGGATG   (-64bp) (+10bp)  (X2)
              CCAAACTTTT
       TTGGCCGTGATGGGCAGTTC         MP              GGCAACTGGGTTGGGAA·······················   ···················CGGGATG    (-59bp) (+16bp)
```

#### Cas9-PAM-out

```
WT  :  CGATCGCCGCAG        GGCATCCAAGTATCGCCATCCGGGATGCGACTGCTCAATGGCCAAC      CTGTGGACGCCAAGG

       CGATCGCCGCAG        GGCATCCAAGTATCGCCATCCGGGATGCGACTGCTCAATGGCCA··      CTGTGGACGCCAAGG              (-2bp)  (+2bp)  (X2)
                                                                     TG
       CGATCGCCGCA·        ····TCCAAGTATCGCCATCCGGGATGCGACTGCTCAATGGC·A··      ··GTGGACGCCAAGG              (-8bp)  (+1bp)
                                                                  G
       CGATCGCCGCAG        GGCATCCAAGTATCGCCATCCGGGATGCGACTGCTCA·········      CTGTGGACGCCAAGG              (-9bp)  (+1bp)
                             G
       CGATCGCCGCA· TCGAT  ·············GCCATCCGGGATGCGACTGCTCAATGGC·A··      ··GTGGACGCCAAGG              (-14bp) (+5bp)
       CGATCGCCGCAG CTGCGATCG ············································   CTGTGGACGCCAAGG              (-47bp) (+9bp)
       CGATCGCCGC·· GGCCAC ··············································      ··GTGGACGCCAAGG              (-50bp) (+6bp)
       CGATCGCCGCAG        GG···················GATGCGACTGCTCAATGGACT·G      CTGTGGACGCCAAGG              (-23bp) (S 3bp)
```

**Cas9-PAM-in**

```
WT  :  TTGGCCGTGATGGGCAGTTCCGG       TGCCGGAAAGACGACCCTGCTGAATGCCCTTGCCTTTCGATCGCCGCAG    GGCATCCAAGTATCGCCATCCGGGATG

       TTGGCCGTGATGGGCAGTTCCGGGAT    .................................................    GGCATCCAAGTATCGCCATCCGGGATG   (-49bp) (+3bp)
```

**Fig S1 - Related to Fig.2 - Variety of resistant alleles produced by the regular Cas9 and nickase versions combined with our CopyCat elements. a-e.** These sequences were recovered by sequencing $F_2$ males from crosses showing super-Mendelian inheritance rates in Fig. 2. The *w2*, *w8* and *w9* gRNA sequences are indicated in blue, orange and green, respectively. PAM sequences are highlighted in red. Dots represent deletions observed in each genotype. Nucleotide insertions were highlighted in pink while substitutions appeared underlined. The different resistant allele categories identified in each genotype are shown. **a.** Large deletions represent mutations erasing more than 50bp and spanning both cut sites. **b.** Resistant alleles contain single mutations occurring at one cut site but not at the other. In all cases, these mutations were smaller than 50 nucleotides. Also, our Sanger sequencing analysis displayed messy traces where the exact sequence could not be determined. These alterations showed multiple peaks (MP) in our Sanger sequencing analysis and suggest that the gRNA could cut after germline formation, causing genetic mosaicism. These events are represented by an MP when needed. **c.** Simultaneous resistant alleles were also formed at both target sites. All these deletions were smaller than 50 base pairs. **d.** Large insertions bigger than 200 nucleotides representing partial HDR events were also found. **e.** A combination of deletions and small insertions together in the same animal were also detected. To keep an appropriate sequence alignment in this category, some insertions were allocated under the main alignment sequence line, right after the nucleotide where it was inserted.
